# Regeneration in sponge *Sycon ciliatum* mimics postlarval development

**DOI:** 10.1101/2020.05.04.061218

**Authors:** Anael Soubigou, Ethan G Ross, Yousef Touhami, Nathan Chrismas, Vengamanaidu Modepalli

**Affiliations:** Sorbonne University, Faculté de Sciences et Ingénierie, Campus Pierre et Marie Curie Jussieu, Paris 75252, France; University of Southampton, School of Ocean and Earth Science, Southampton, SO17 1BJ, United Kingdom; Marine Biological Association of the UK, The Laboratory, Citadel Hill, Plymouth PL1 2PB, United Kingdom

## Abstract

Somatic cells dissociated from an adult sponge can re-organize and develop into a functional juvenile. However, the extent to which regeneration recapitulates embryonic developmental signaling pathways has remained enigmatic for more than a century. To this end, we have standardized and established a sponge *Sycon ciliatum* regeneration protocol to achieve consistent regeneration in cell culture. From the morphological analysis, we demonstrated that dissociated sponge cells follow a series of morphological events resembling embryonic and postlarval development. Hence, we propose that sponge regeneration represents somatic development. To support our hypothesis, we performed high-throughput sequencing on regenerating samples and compared the data with regular embryonic and postlarval development of *Sycon ciliatum*. Our comparative transcriptomic analysis illuminates that sponge regeneration is equally as dynamic as embryogenesis. We find that sponge regeneration is orchestrated by complex regulatory mechanisms by recruiting signaling pathways like those utilized in embryonic development to organize into a functional juvenile. In the current study, we lay down the basic framework to study *Sycon ciliatum* regeneration. Since sponges are likely to be the first branch of extant multicellular animal and the sister lineage to nearly all animals, we suggest that this system can be explored to study the genetic features underlying the evolution of multicellularity and regeneration.

## Introduction

The gradient of regeneration widely differs in the animal kingdom and among them, the sponge is an exceptional organism with an extraordinary regenerative ability. Anatomical plasticity and cell differentiation is a characteristic feature of sponges, and many sponge cells are capable of transdifferentiation [1, 2]. Dissociated cells from an adult sponge have the unique ability to reaggregate and fully reconstitute into a functional sponge [3-5]. Despite its significance, we still lack an in-depth understanding of the molecular mechanisms and cellular basis of sponge regeneration. Sponges are the ancient extant branch of metazoans (branching order is still debated) [6] and understanding their cellular behavior or morphogenesis during regeneration can provide insights into the evolution of multicellularity and regeneration. Calcareous sponges have been extensively studied in the past centuries and analysis of their development has significantly influenced evolutionary theory. Terms such as gastrulation and metamorphosis were first coined in syconoid species from the classic Calcaronea studies [7]. *Sycon ciliatum*, a calcareous sponge, is widely explored in embryonic and regeneration studies [8, 9].

In *S. ciliatum*, embryogenesis is viviparous. In brief, after fertilization the embryos through early cleavages develop into a rhomboid shape. Subsequent cell divisions result in the formation of a cup-shaped embryo (stomoblastula). The embryo is composed of granular macromeres adjacent to the choanocytes and ciliated micromeres pointing into the embryonic cavity. After inversion, the cilia are positioned on the outer surface of the larva and eventually the ciliated amphiblastula larva migrates into the radial chamber and swims out of its parent **(Fig.1)** [8-10]. The amphiblastula larva is composed of a single layer of embryonic cells: ciliated micromeres on the anterior pole, and non-ciliated granular macromeres on the posterior pole. During metamorphosis, the ciliated cells of the anterior half undergo an epithelial-to-mesenchymal transition and form the inner cell mass; the choanocytes will differentiate from these cells. The macromeres envelop the inner cell mass and gradually transform into pinacocytes (pinacoderm). After 24 hours following settlement, the spicules (monaxons) are produced by sclerocytes which have differentiated from the inner cell mass. A choanocyte chamber is formed in the inner cell mass and develops into the choanoderm. An osculum is formed at the apical end and the juvenile is elongated along the apical-basal axis rises into a syconoid body plan [8, 10] **(Fig.1)**.

**Figure 1:**
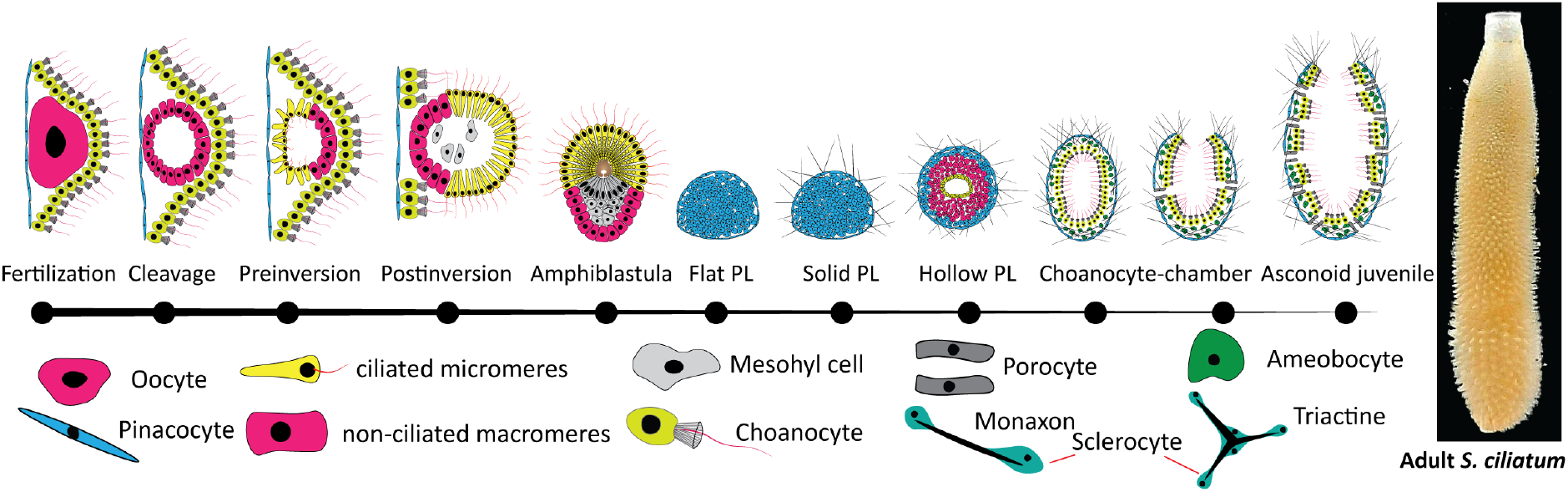
Schematic representation of embryonic and postlarval development stages in *Sycon ciliatum*: depicted based on Leininger, S., et al. 2014, Eerkes-Medrano, D. I. and S. P. Leys 2006. From left to right: oocyte, cleavage, pre-inversion, post-inversion, amphiblastula (swimming larva), flat post larva, solid post larva, hollow post larvae, and choanocyte chamber and asconoid juvenile.

Many studies have been dedicated to *S. ciliatum* regeneration to understand their morphological signatures, starting from Wilson 1907 [3], the pioneer of this topic. Despite their strong morphological resemblance to embryonic and postlarval development, to date, no study has systematically compared them. Several transcriptome studies of regeneration in cnidarians and bilaterians have demonstrated that many developmental pathways are recruited in regeneration [11-14]. Hence we aim to study the regeneration of *S. ciliatum* and compare it to embryonic and postlarval development at both morphological and transcriptional levels. We first carried out *S. ciliatum* regeneration to acquire morphological data for comparison to regular embryonic and postlarval development and classified the regeneration into seven key stages. Then, we collected the regenerating aggregates at defined time points and performed high-throughput sequencing to compare the transcriptional profile with those of normal embryonic and postlarval development.

## Results and Discussion

### *Sycon ciliatum* regeneration morphologically resembles embryonic and postlarval development

First, we standardized and established a *S. ciliatum* regeneration protocol to achieve consistent sponge regeneration in cell culture. Dissociated cells sink to the bottom of the dishes within 1 hr of dissociation. Among these cells, we detected choanocytes, amoeboid cells, pieces of spicules and round cells. The majority of the cells lose their cell morphology. In 4-6 hours post dissociation (hpd), loose multicellular aggregates of diverse shapes (∼10 µm) are formed; they are known as primmorphs. The contacts between the cells were not strong, and upon shaking the aggregates fell apart into separated cells. Within 24 hpd, these primmorphs significantly increased in size (∼100 µm) predominantly due to the incorporation of cells from suspension or by fusing with other aggregates **(Fig.2 A-C)**. The primmorphs have no defined surface epithelium; on the surface pinacocytes and amoebocytes, as well as flagella and collars of choanocytes were noticeable and similar types of cells can be found inside the primmorphs **(Supplementary file 1-3)**. During the early stages the cells are actively reorganized and at 48 hpd most of these cells have lost their typical features, primarily due to transdifferentiation **(Fig.2 J-K)**. The primmorphs become denser and spherical and two layers of cell types can be identified: a uniform single-cell layer of pinacocytes on the surface and loosely packed granular cells inside the primmorphs **(Fig.2 D-F) (Supplementary file 3)**. Earlier studies coined this phenomenon of forming bilayer cells as redevelopment, or somatic development [5, 15]. Choanocytes on the surface were oriented with their flagella outwards and retained their characteristic features up to 2-3 days post dissociation (dpd) and after that, they started gradually losing their typical features (flagellum and microvilli) and becoming indistinguishable from other cells **(Fig.2 I, L)**. Largely, two types of primmorphs were observed in the culture: the first one is granulated cell packed spheroid with a small bubble and another one is with a large bubble and a small inner cell mass (termed as blow-outs by Julian S. Huxley. [4]), an interesting morphology that resembles a blastula with blastocoel. Eventually, a single continuous cavity is formed between the external single cell layer of pinacocytes and internal cell mass **(Fig.2 H)**. At 3 dpd, the primmorphs were entirely covered with pinacocytes, which may have possibly formed through transdifferentiation from choanocytes **(Fig.2 J-L)**.

**Figure 2:**
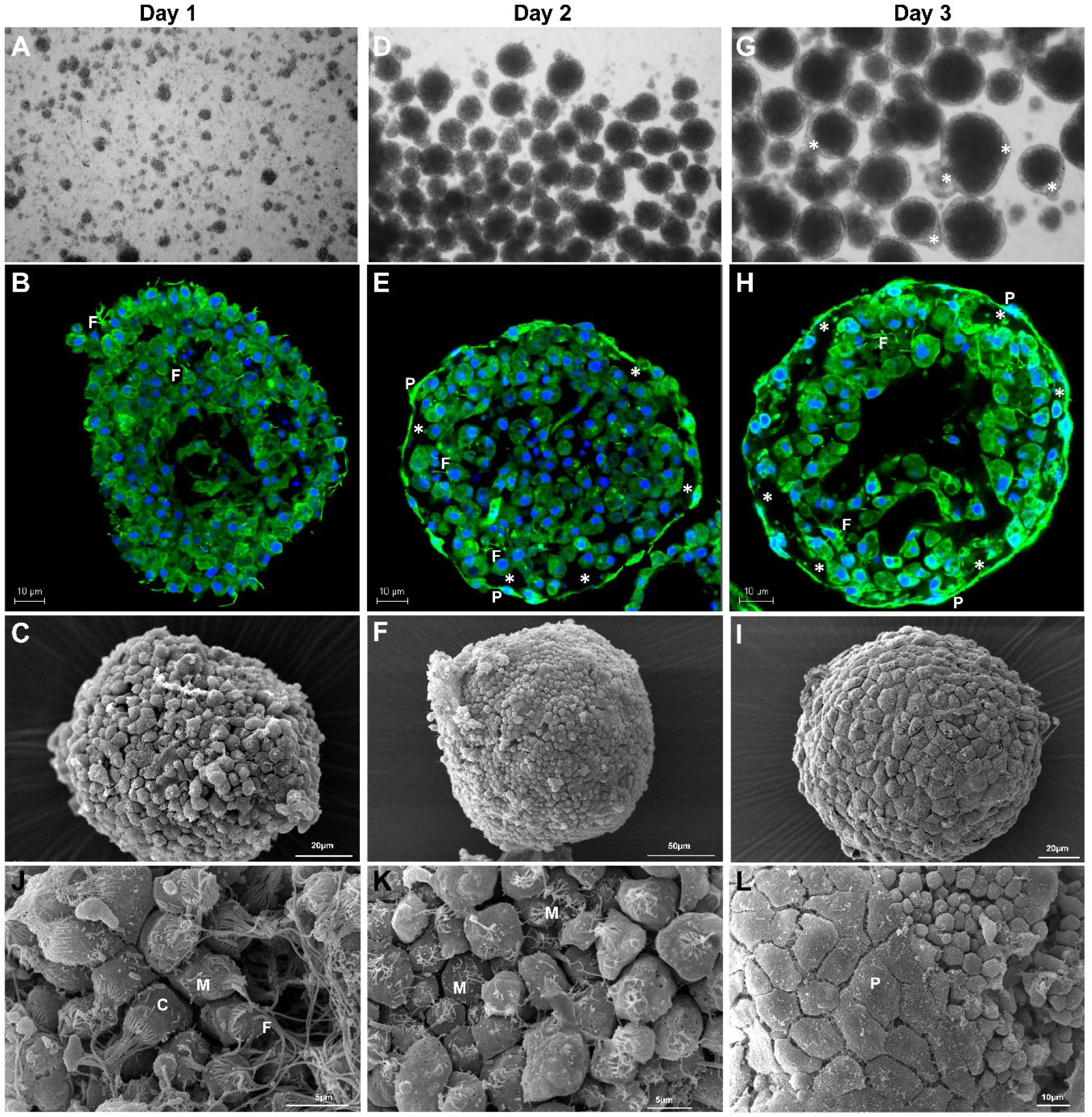
Morphological and cellular events during the primmorphs stage. The regeneration proceeds with the aggregation of dissociated cells and transforming the loosely connected external cells to an intact single layer of pinacocytes. **(A-C)** Initial multicellular aggregates in the culture (24 hpd), loosely aggregated primmorphs, the primmorphs have no defined surface epithelium and made of diverse cell types. **(D-F)** A general view of primmorphs in the culture after 2 dpd, **(E & F)** an intact single layer of cells surrounding primmorphs, separating from inner cell mass, similar to blastocyst. **(G & H)** The external ameboid or cuboidal cells transformed into flat pinacocytes, the external cavity (*) resembles the extraembryonic cavity with a single layer of flat cells. **(E, F, H, I)** Fusing the external cells surrounding the primmorphs, accompanied by the transdifferentiation of the choanocytes and other external ameboid cells. **(J-L)** choanocytes transdifferentiation into pinacocytes: **(J)** choanocytes with intact flagella and microvilli, **(K & L)** gradually these morphological features were lost through transdifferentiation and forms falt later of pinacocytes. Images are taken with **(A, D, G)** bright field, **(B, E, H)** confocal, **(C, F, I)** SEM microscope. P: pinacocyte; C: choanocytes; F: flagella; M: microvilli; (*): external cavity.

Around 5 dpd spicules start appearing from the primmorphs and simultaneously multiple ciliated chambers start appearing inside the cell mass. Similarly to regular postlarval development [8], the monaxons were formed first **(Fig.3 A, D)**. Over the next 6-10 days, the ciliated chambers expanded and became lined with choanocyte epithelium **(Fig.3 G-I) (Supplementary file 4 & 5)**. Gradually, these choanocyte chambers fuse into a single cavity lined with choanocytes (also known as choanoderm). Around 16 dpd a single extended cavity (spongocoel) with osculum forms at the apical end, and simultaneously multiple porocytes with ostia are formed **(Fig.4 A, C) (Supplementary file 6)**. These juveniles continue elongating along the apical-basal axis, and long straight spicules form a crown around the osculum. Further development can be described as an increase in the complexity of spongocoel and the development of intermediate porocytes with ostia **(Fig.4 E-J) (Supplementary file 7)**.

**Figure 3:**
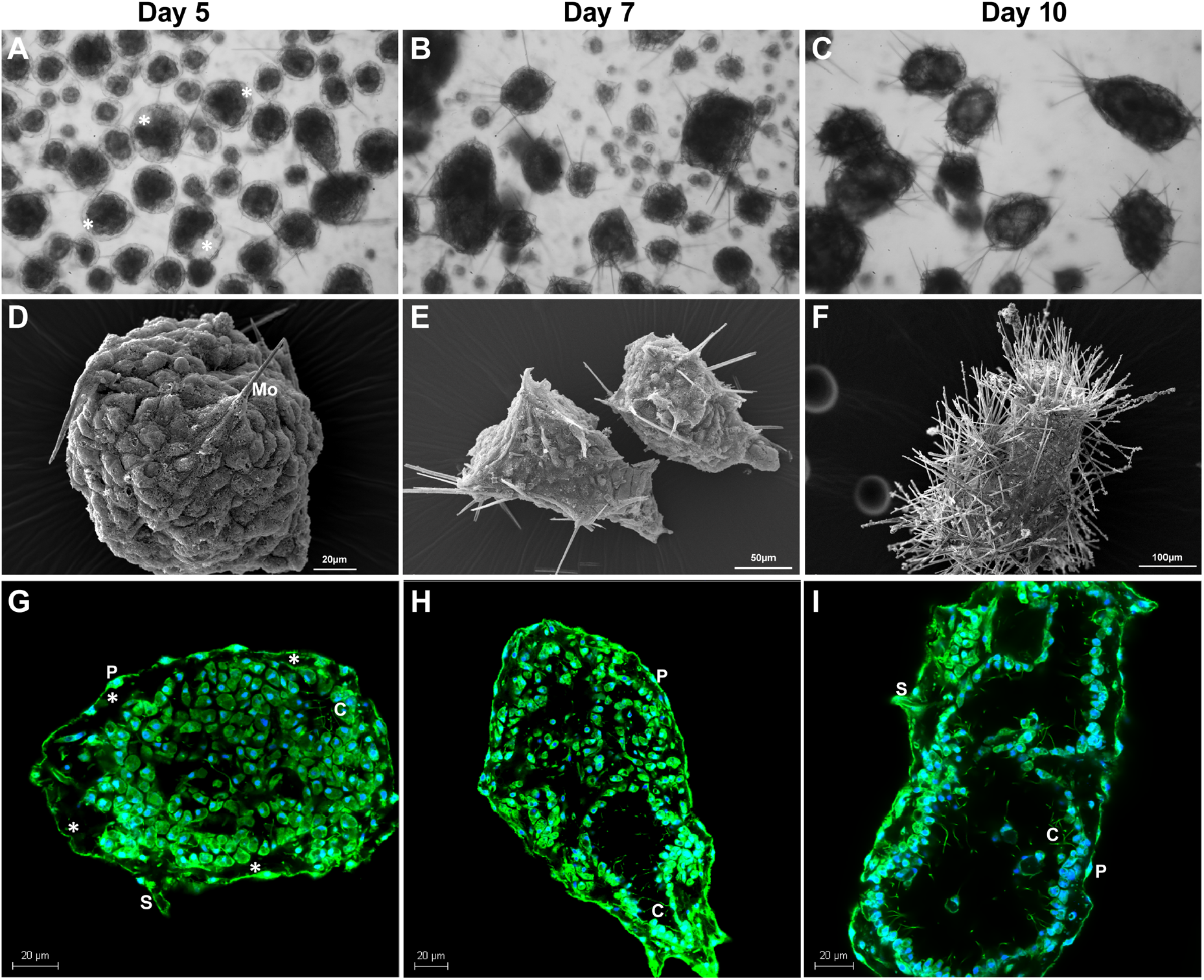
Specule formation during regeneration. **(A-F)** general view of primmorphs developing spicules (4-10 dpd). **(D & G)** Primmorphs developing multiple chambers surrounded by ciliated cells, simultaneously the spicules start appearing as monaxons. **(G-I)** expansion and integration of choanocyte chambers and eventually developing into choanoderm with a single layer of choanocytes surrounding the atrium. **(F & I)** note that ostium and osculum are not formed yet. Images are taken with **(A, B, C)** bright field, **(D, E, F)** SEM microscope, **(G, H, I)** confocal. C: choanocytes; S: sclerocytes; P: pinacocyte; Mo: monaxons. (*): external cavity.

**Figure 4:**
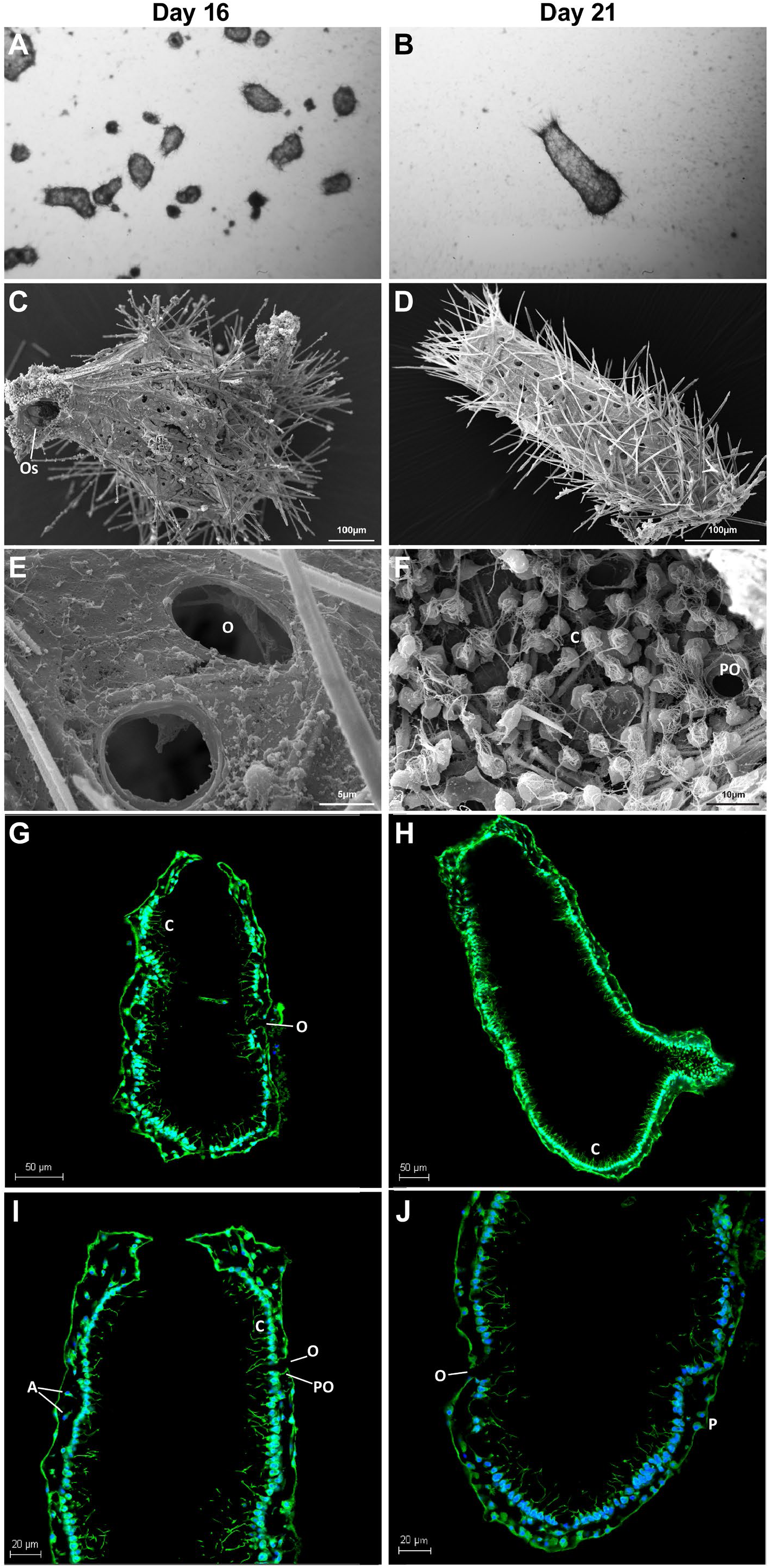
Developing a functional juvenile. **(A, B)** General view of pre-juvenile and juvenile stages. **(C-J)** Osculum opening at apical end and multiple porocytes form ostia. **(C)** The appearance of ostium and osculum is synchronized. **(G)** Ostia in the regenerated juvenile, **(H)** a view of ostia from inside, ostia is surrounded by choanocytes. O: Ostia; PO: porocytes; C: choanocytes; S: sclerocytes ; P: pinacocyte; Os: osculum; A: archaeocytes or amoebocytes.

Regeneration in *S. ciliatum* displayed a multitude of complex morphogenesis steps, progressing through a series of aggregation, organization, and somatic-development events. Based on our current observations and multiple pieces of evidence from previous studies [3-5], we classified sponge regeneration into seven stages based on the morphological and cellular events: I) Aggregation, II) Primmorphs, III) Primmorphs with monaxons, IV) Primmorphs with ciliated chambers, V) Choanoderm, VI) Pre-Juvenile (Ascon stage), VII) Juvenile; the details are presented in the following table **(Table 1, 3**^**rd**^ **column)**. Remarkably these morphological signatures are vastly comparable to embryonic and postlarval development of *S. ciliatum* **(Fig.1 and Table 1, 4**^**th**^ **column)** [1, 8, 9]. To gain further insights into their morphological overlap, we compared regeneration with both embryonic and postlarval development. The embryogenesis of calcaronean sponges was previously well described on light and electron microscopy levels [1, 8, 9]. The morphological comparison was performed using the data from the published *S. ciliatum* embryonic development studies (Leininger, S., et al. 2014 and Eerkes-Medrano, D. I. and S. P. Leys 2006). As demonstrated in **Table 1**, apart from initial stages of primmorphs at day 1, 2 and 3, the majority of *S. ciliatum* regeneration steps are overlapping with a normal *S. ciliatum* development directly after metamorphosis including S2, S3, S4, S5 and YS [8].

**Table 1:**
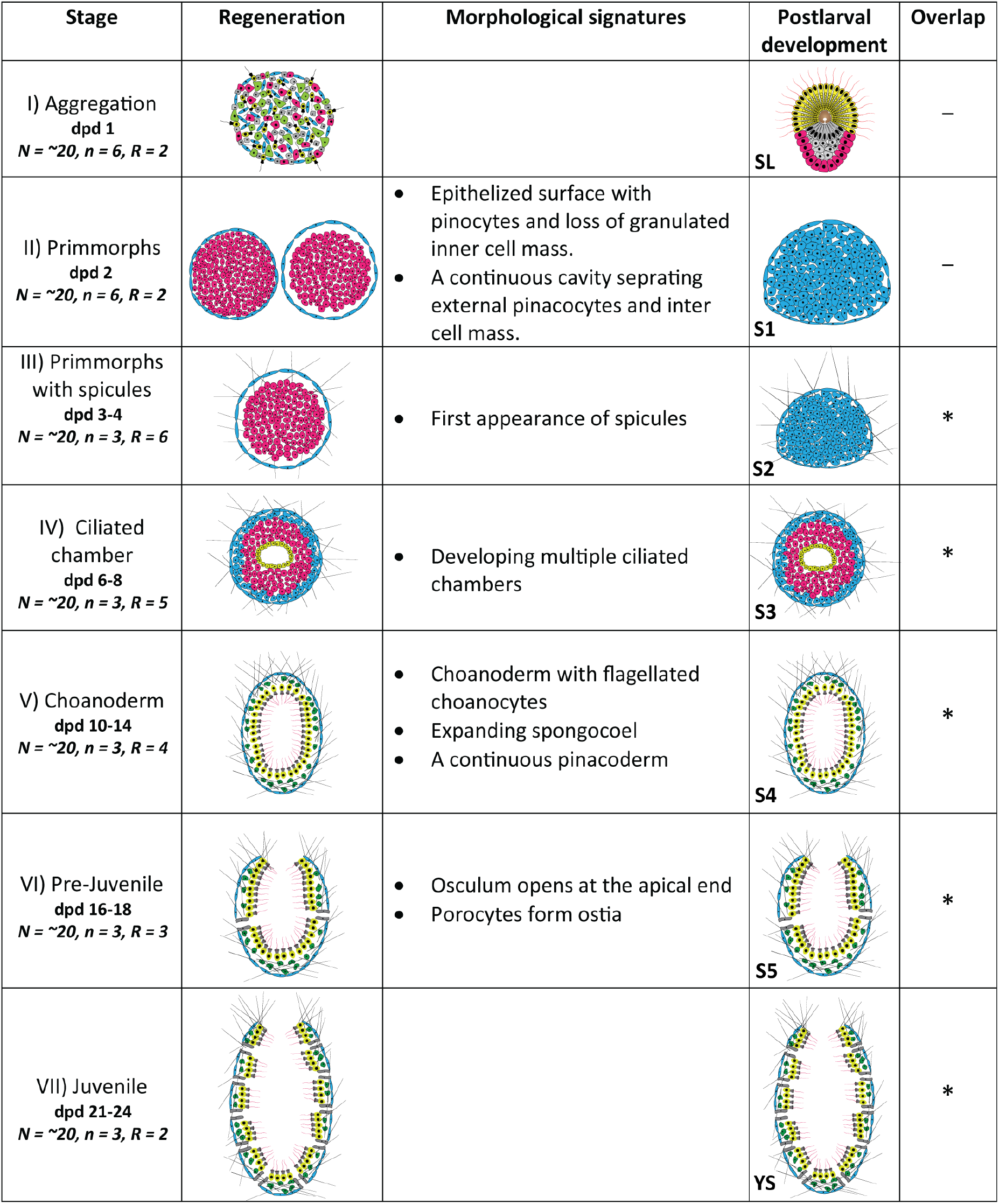
Summary of the morphological and cellular events during regeneration in *S*.*ciliatum*. The table details various regeneration steps and their overlaps with postlarval development. 4^th^ column (postlarval development) depicts the morphological events at postlarval development of *S. ciliatum*. 1^st^ column details number of samples collected for RNA-sequencing at each stage of regeneration, N=number of pooled regenerating structures per well, n=number of pooled regenerating structures per replicate, R=number of replicates per stage.

| Stage | Regeneration | Morphological signatures | Postlarval development | Overlap |
| --- | --- | --- | --- | --- |
| I) Aggregation<br>dpd 1<br>$N = \sim 20, n = 6, R = 2$ | | | SL | — |
| II) Primmorphs<br>dpd 2<br>$N = \sim 20, n = 6, R = 2$ | | Epithelized surface with pinocytes and loss of granulated inner cell mass.<br>A continuous cavity separating external pinacocytes and inter cell mass. | S1 | — |
| III) Primmorphs with spicules<br>dpd 3-4<br>$N = \sim 20, n = 3, R = 6$ | | First appearance of spicules | S2 | * |
| IV) Ciliated chamber<br>dpd 6-8<br>$N = \sim 20, n = 3, R = 5$ | | Developing multiple ciliated chambers | S3 | * |
| V) Choanoderm<br>dpd 10-14<br>$N = \sim 20, n = 3, R = 4$ | | Choanoderm with flagellated choanocytes<br>Expanding spongocoel<br>A continuous pinacoderm | S4 | * |
| VI) Pre-Juvenile<br>dpd 16-18<br>$N = \sim 20, n = 3, R = 3$ | | Osculum opens at the apical end<br>Porocytes form ostia | S5 | * |
| VII) Juvenile<br>dpd 21-24<br>$N = \sim 20, n = 3, R = 2$ | | | YS | * |

### The gene expression of regenerating *Sycon ciliatum* is as dynamic as postlarval development

A long-standing question in the field of sponge regeneration is whether and to what degree regeneration recapitulate embryonic developmental pathways. Extensive morphological similarities between regeneration and postlarval development (PLD) have driven us to explore the gene expression pattern of both processes. We sequenced regenerating samples at 7-time points spanning from 1 to 24 days including dpd 1, 2, 3-4, 6-8, 10-14, 16-18 (PJUV) and 21-24 (JUV). For postlarval development, the RNAseq datasets spanning 7-time points were collected from previously published data (PRJEB7138 & PRJEB5970) including amphiblastula (swimming larva), flat post larva (S1), solid post larva (S2), hollow post larvae (S3), choanocyte chamber (S4 & S5) and asconoid juvenile (PY) stages [8]. For comparative analysis, we processed both data sets through similar pipelines as described in the material and methods section. To assess the global transcriptomic profile underlying postlarval development and regeneration, we performed principal component analysis (PCA) on individual data sets. We found that both regeneration and PLD follow similar PCA distribution and surprisingly the transcriptional changes during regeneration are as variant as PLD **(Fig.5 A&B)**. We also performed hierarchical clustering (HC) on both datasets to define the major clusters and we identified two major clusters that we classified as early and late stages **(Fig.5 C&D)**. From PCA and HC analysis we observed that clusters in regeneration are organized in a hierarchy consistent with what we observed in the PLD.

**Figure 5:**
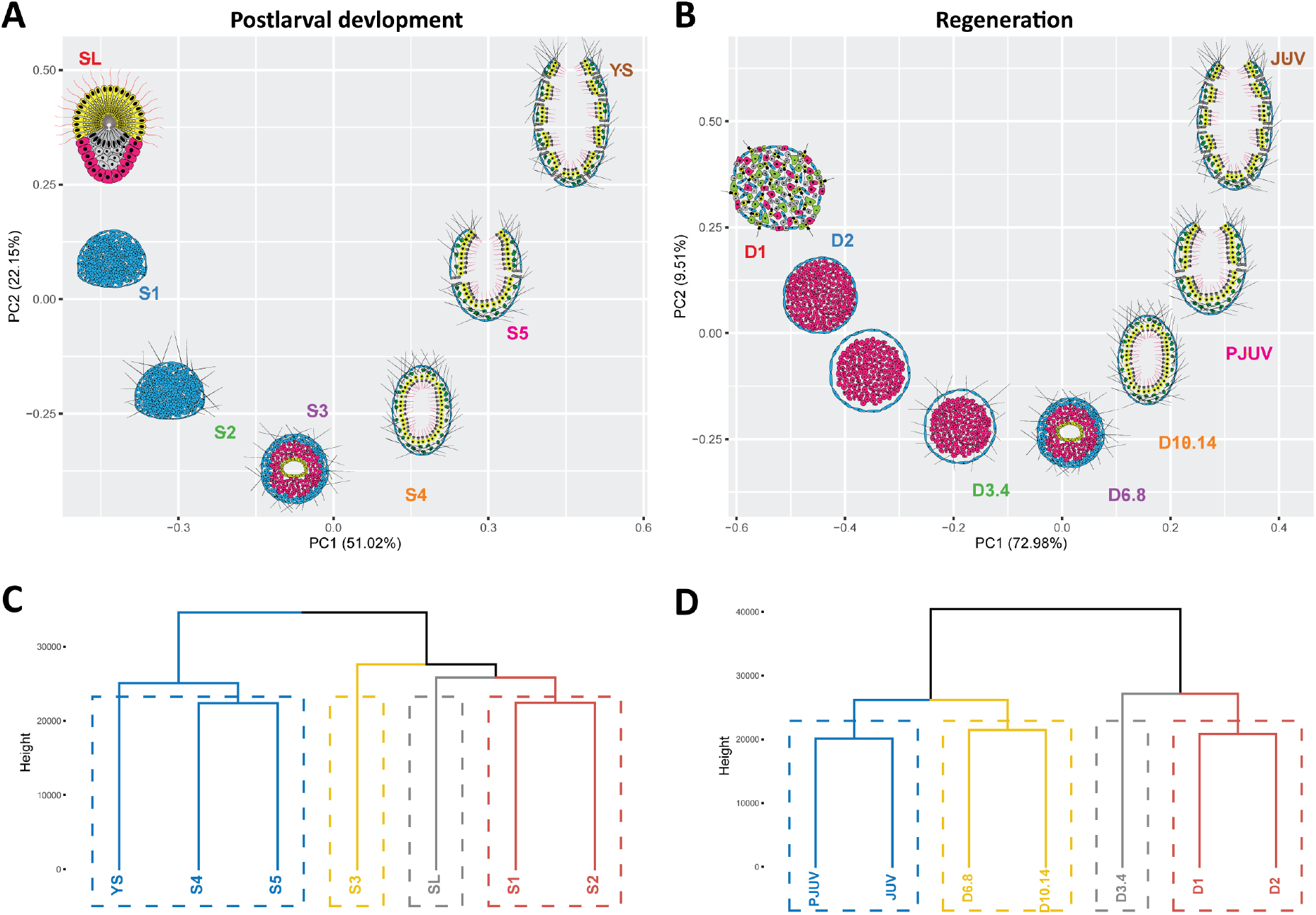
A global overview of postlarval and regeneration transcriptomes. **(A)** Principal component analysis (PCA) of postlarval datasets, samples at SL, S1, S2, S3, S4, S5, and YS stages. **(B)** PCA of regeneration dataset, samples at dpd 1, 2, 3-4, 6-8, 10-14, 16-18 (PJUV) and 21-24 (JUV). **(C & D)** hierarchically clustering (HC) on postlarval development **(C)** and regeneration **(D)** datasets revealed two major clusters that we classified as early and late stages. This analysis confirmed that clusters in regeneration are organized in a hierarchy consistent with what we observed in the PLD.

We then directly compared the transcriptomic variation among regeneration and PLD using PCA. Despite their transcriptomic dynamics, regeneration and PLD fall apart from each other mainly at PC1 proportion. However, the profiles of regeneration and PLD began to converge as they progress through development **(Fig.6 A)**. This indicates that the transcriptomic dynamics of regeneration and PLD are probably similar at advanced stages. Similar conclusions were also drawn in HC analysis: the late stages (D10-14, PJUV & JUV) from regeneration have clustered with S5 and PY from PLD **(Fig.6 B&C)**, further suggesting that the later stages of both regeneration and PLD may express genes with maximum correlation. High variance among early time points of regenerating and developmental samples indicates an alternative path was taken during the early stages of regeneration. As noted from the morphology, early stages of regeneration are not strictly a morphological recapitulation of development with potential for extensive temporal plasticity during early regeneration.

**Figure 6:**
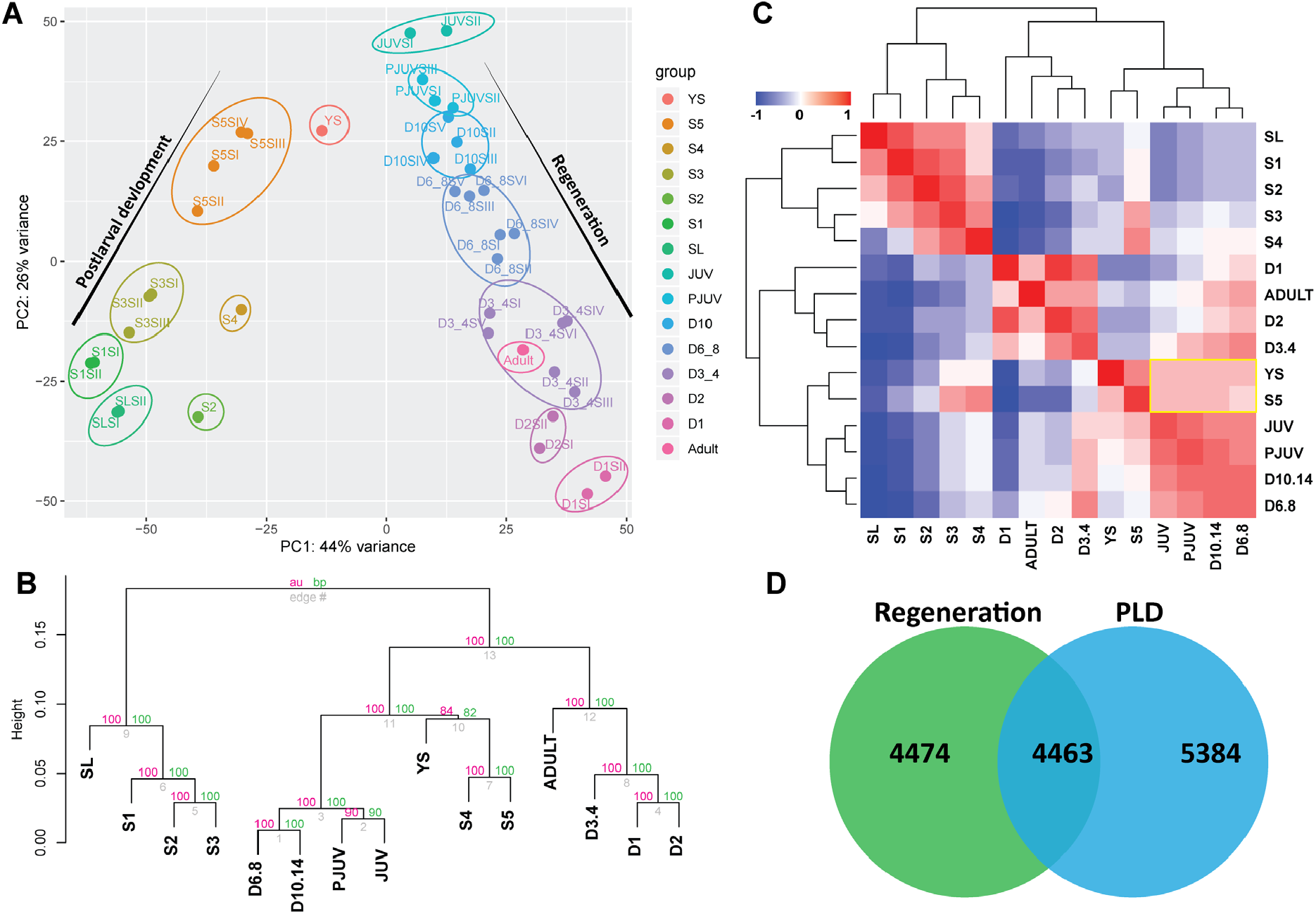
Global transcriptomic comparison of the regeneration and PLD samples. **(A)** PCA of larval development versus regeneration datasets. Regeneration exhibits similar transcriptomic variation as embryogenesis. The regeneration samples collected at D10-14, PJUV and JUV stages are close to the S5 and PY stages of PLD. **(B)** Hierarchical clustering, the dendrogram of both datasets obtained with pvclust is shown with the bootstrap values on each node. The dendrogram shows a significant correlation among late stages of regeneration (D10-14, PJUV and JUV) with S5 and PY stages of PLD. **(C)** Comparison of differentially expressed genes between PLD and regeneration datasets, similarity matrix for significantly differentially expressed genes (padj (FDR) < 0.01 for any timepoint comparison against Day 1 for regeneration and S1 for PLD). The yellow box highlighting the similarity between late stages of regeneration (D10-14, PJUV and JUV) with S5 and PY stages of PLD. **(D)** The number of differentially expressed genes in regeneration is highly comparable to PLD and ∼50% of genes are commonly expressed.

A further comparison was carried out on the significantly differentially expressed genes (padj (FDR) < 0.01) at any time point compared to day1 for regeneration and S1 for PLD.

Regeneration exhibited 8937 differentially expressed genes (DEG), which is comparable to 9847 DEG in PLD, suggesting that regeneration exhibits dynamic gene expression equivalent to PLD. Of these dynamically expressed genes, approximately 50% of genes (4463) are commonly expressed in both data sets **(Fig.6 D)**, suggesting that half of the PLD genes were deployed during regeneration. As regeneration employs 50% of PLD genes, we intended to compare the expression profiles of these commonly differentially expressed genes. We subjected the candidates to Fuzzy c-means clustering [16] to group the genes based on the expression profile **(Fig.7 A&B)**. The clustered genes were compared between regeneration and postlarval development. We found that the majority of the regeneration gene clusters exhibited significant overlap with specific PLD clusters **(Fig.7 C&D)**. We conclude that regeneration partly resembles PLD by displaying similar gene expression profiles.

**Figure 7:**
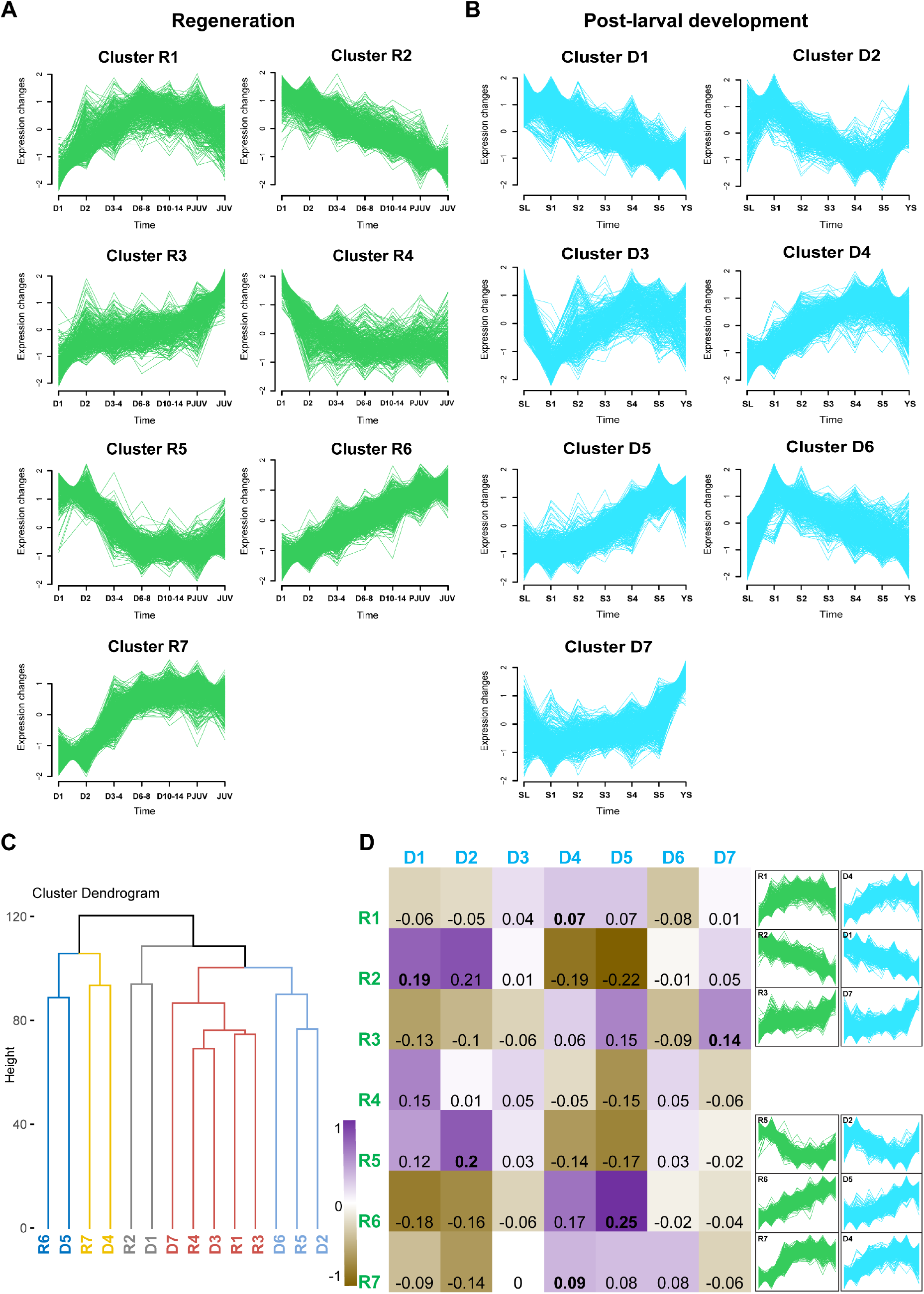
Regeneration gene clusters overlapping with PLD gene clusters. A total of 4468 commonly expressed genes between regeneration **(A)** and PLD **(B)** are subjected to Fuzzy c-means clustering. Based on their gene expression profile we generated 7 clusters in both datasets. **(C)** Hierarchical clustering. The dendrogram of all gene expression clusters from regeneration and PLD. Majority of regeneration gene modules clustered with PLD gene modules. **(D)** Cluster overlap table of regeneration versus PLD gene expression clusters produced via Pearson correlation analysis. Bold text indicates the best cluster overlap and their matching gene expression profiles are presented on the right side of the table.

Overall, we find that late stages from both regeneration and PLD have a high correlation, thus the major variation in these data is among the early stages. Since the regeneration is not starting from a fertilized single-cell embryo or amphiblastula larva, early regeneration stages do not necessarily need to follow a similar pattern of gene expression profile to normal embryonic and PLD development. Despite its rudimentary embryonic and swimming larval morphology, the early primmorphs were not close to normal swimming larva in gene expression profile, however the subsequent stages do resemble postlarval development **(Fig.6)**. The early primmorphs are unique, as observed from morphological analysis, and multiple cellular events at this stage are expected to have unique gene expression.

### Analyzing *S. ciliatum* regeneration gene expression

To the best of our knowledge, sponge regeneration from dissociated somatic cells has not been examined at the transcriptome level. We analyzed the regeneration gene expression data to identify developmental and multicellular regulatory genes. We performed a GO-term enrichment analysis to reveal gene network acting during regeneration. The analysis revealed that a set of transcription factors tightly controls the regeneration process. A broad range of eumetazoan developmental regulatory genes were expressed during regeneration, gene families such as *Wnt, Tgfb, Fzd*, and *Smad* were differentially expressed **(Fig.8 A)**. Surprisingly, the majority of listed genes were previously identified in *S. ciliatum* during embryonic and postlarval development [8, 17-19]. From the pathway enrichment analysis we identified Wnt, TGF-beta, Notch, Hedgehog, FGF and EGF signaling pathways, and the majority of core pathway components were detected **(Fig.8 A, Supplementary file 8)**. Each of these signaling pathways triggers several downstream signaling cascades regulating cell behavior such as cell-cell and cell-ECM interactions [20]. Along with other biological processes, apoptosis plays an important role during regeneration [21, 22]. Apoptosis mediated regeneration permits cell elimination and reorganizes cell composition. Major cell death or apoptotic associated pathways including Wnt signaling, oxidative stress response and p38 MAPK pathway were expressed during *S. ciliatum* regeneration **(Supplementary file 8)**, revealing the importance of apoptosis in remodeling the primmorphs to initiate re-development.

**Figure 8:**
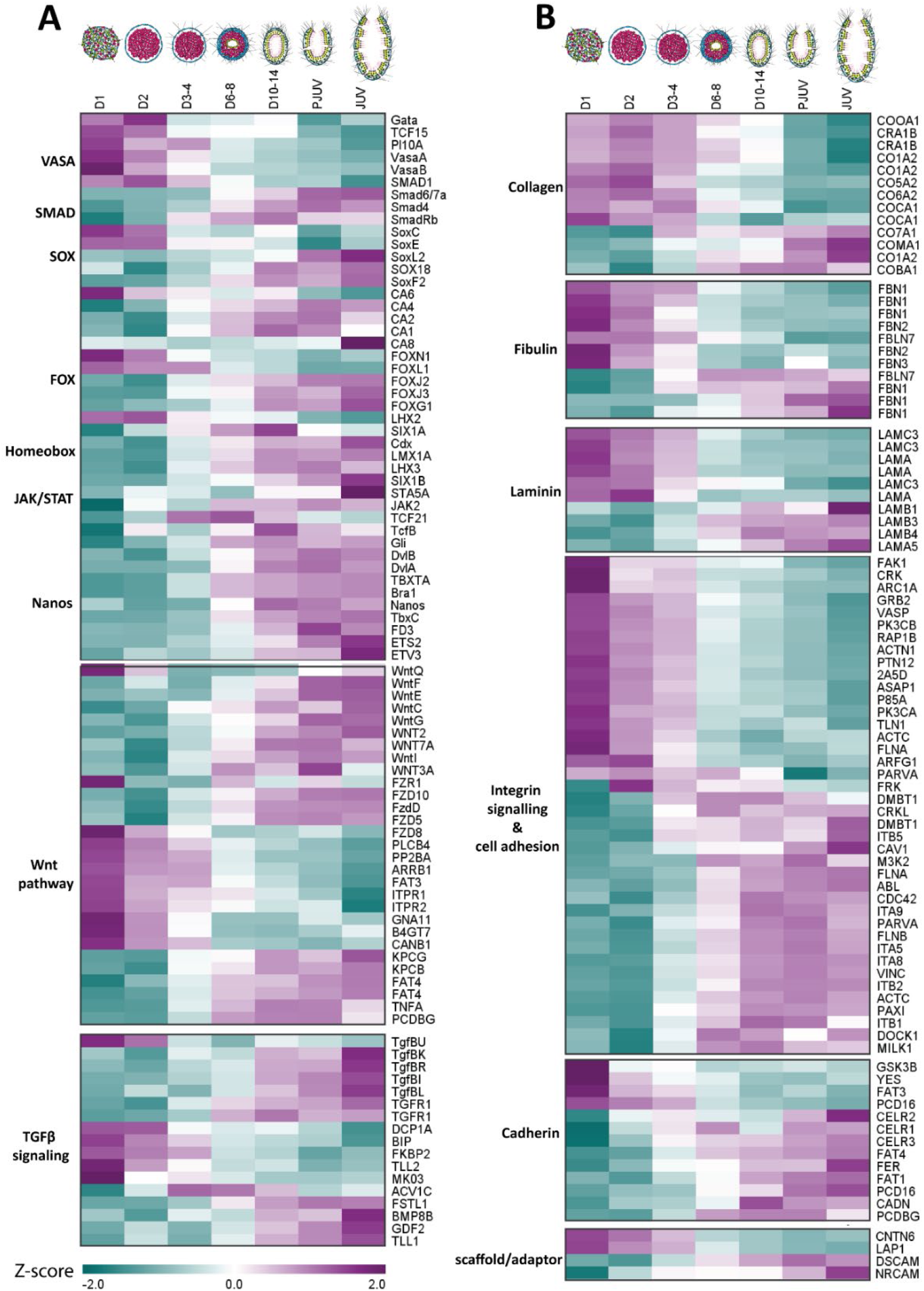
Gene expression across *S. ciliatum* regeneration. Heatmap of significantly DEG. **(A)** A heatmap displaying a selected list of transcriptional factors and Wnt and Tgfb pathways genes. The majority of listed genes are well known to associate with a range of eumetazoan developmental regulatory pathways and were previously identified in *Sycon ciliatum* embryonic development. **(B)** A heatmap showing relative expression of ECM and cell adhesion and motility (integrin, cadherin, and scaffold) genes, a broad range of genes started expressing within 24 hpd, possibly involved in cell-aggregation, adhesion, and motility during regeneration. Heatmap of scaled expression (row z-score) of integrins with significantly different expression (padj (FDR) < 0.01 for any timepoint comparison against Day 1).

After tissue dissociation, the initial cellular source constitutes a heterogeneous pool of adult somatic cells. During the initial regeneration phase (day 1 to 3) the dissociated cells re-aggregate and organize into a morphologically distinguishable structure **(Fig.2)**. During the development of multicellular organisms, cells change their position extensively and this process is governed by cell-adhesion and motility molecules. Similarly, the sponge regeneration is guided through a series of cellular events including initial cell aggregation and followed by cell organization. Hence, we analyzed the genes associated with cell adhesion and cell mobility molecules. Using GO term analyses of DEG, we identified members associated with cell-adhesion gene families, including those that encode cell-surface receptors, cytoplasmic linkers, and extracellular-matrix proteins **(Fig.8 B)**. Expression of these cell-cell and cell-ECM adhesions signify highly integrated networks crucial for the individual cells to adhere and organize into a multicellular structure [23]. Cadherins mediate cell-cell adhesion broadly regulating morphological aspects during development [24, 25]. Several cadherin pathway associated genes were significantly differentially expressed **(Fig.8 B)**. Along with cadherins we also identified integrin pathway associated genes. Cell migration is a process that is highly dependent on adhesion and junction molecules such as integrins [26, 27]. The co-expression of cadherin, integrins and other down-stream scaffolding and adaptor proteins such as RhoGTPases, tyrosine kinases, and phosphatases, suggests that together these molecules contribute to initiating the early events of regeneration. Integrins mediate contact between cells and ECM in many organisms by binding to ECM proteins [28-32]. Three major extracellular matrix structural proteins fibronectin, collagen and laminin displayed differential expression **(Fig.8 B)**, demonstrating the importance of ECM molecules during regeneration. In summary, our analysis indicates that initial phases of regeneration, including cell aggregation and organization, are guided by a set of cell-cell and cell-ECM adhesion associated molecules.

### Lessons to learn on the origin of multicellularity over the sponge regeneration model

The factors that have driven the emergence of multicellular animals from their unicellular ancestors have yet to be addressed. Several studies on unicellular relatives of animals including choanoflagellates, filastereans, and ichthyosporeans have significantly contributed to understanding how unicellular organisms transform into multicellular structures such as colonial choanoflagellate [33-35]. This suggests similar transitions may have taken place in the animal ancestors, which eventually evolved into a stable and functional multicellular animal [36, 37]. The current hypothesis on the origin of multicellular animals generally agrees that choanoflagellates share a common ancestor with animals **(Fig.9)**. This idea is based on the molecular phylogeny and morphological similarity between choanoflagellates and sponge choanocytes **(Fig.9 A-C)** [38]. Further, the choanoflagellate colonies partially resemble the choanocyte chambers of the sponge [37, 39, 40]. In addition to their unicellular sister lineages, we also need to investigate other close relatives of the first multicellular animals.

**Figure 9:**
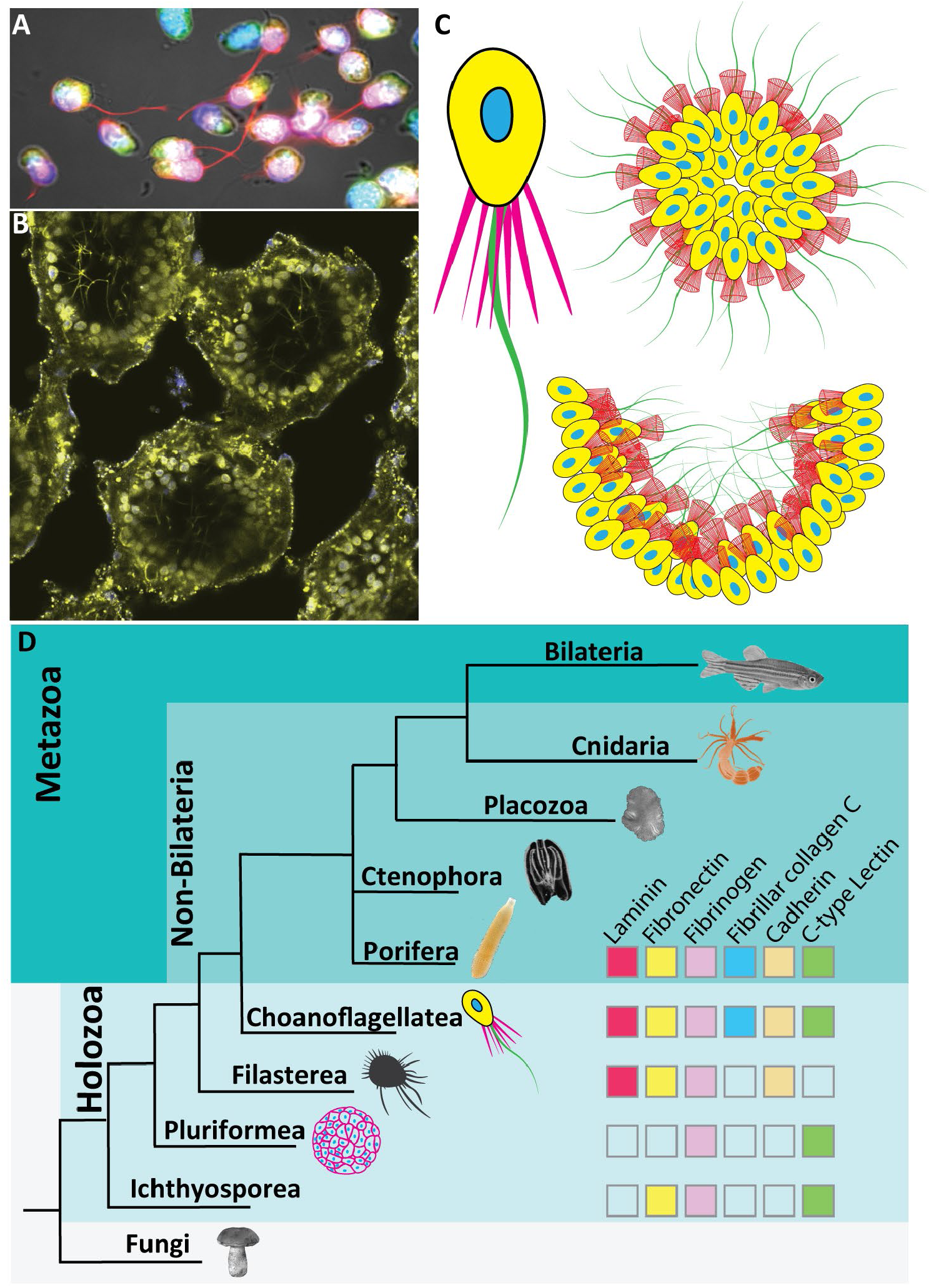
The origin of animal multicellularity. **(A-C)** Morphological similarities between choanocytes and choanoflagellates. **(A)** *S. ciliatum* choanocytes among dissociated cells; **(B)** choanocyte chambers of sponge *S. ciliatum*; **(C)** unicellular and multicellular (colonial) forms of choanoflagellates. **(D)** Phylogenetic tree presenting animals and closely related unicellular holozoan lineages. The position of Ctenophora and Porifera is indicated as a polytomy due to current uncertainty on their evolutionary origin. Animal ECM and cell-adhesion family proteins found in unicellular holozoan lineages; empty boxes indicate absence.

Studying the early regeneration events in sponge, including the signalling molecules triggering the aggregation of cells, can provide a window into understanding the transition from unicellular to multicellular life forms. Recent studies on multiple unicellular holozoans revealed several genes related to multicellularity including cell differentiation, cell-cell, and cell-ECM adhesion **(Fig.9 D)** [41-44], suggesting these factors have evolved before the origin of multicellular animals. Indeed, in our current analysis on *S. ciliatum* regeneration gene expression, we found that several cell signalling and adhesion protein families including cadherin, integrins, and ECM molecules were highly expressed within 24 hrs from dissociation **(Fig.8 B)**. The main focus of the current study is to test the homology between regeneration and embryonic development, hence sampling was prioritized based on morphology with the first sample collected 24 hr following dissociation. To gain an in-depth understanding of the initial signalling cascades triggering the aggregation a tight sampling during the initial aggregation stages would be beneficial. Future studies in this direction will provide further insights into the evolutionary mechanisms underlying the origin of multicellular animals.

## Conclusions

Sponge regeneration is a powerful system to understand the cellular and molecular mechanisms governing animal regeneration in general. This is crucial for understanding the evolution of multicellularity. We compared regeneration and embryonic development in *S. ciliatum* by applying both morphological and RNA-seq analyses to reveal the correlation among these phenomenons. Here we demonstrated that *S. ciliatum* regeneration is governed by core components of the Wnt, Tgfβ, Notch and Hedgehog signaling pathways, which are known to guide various aspects of embryonic development and morphogenesis in extant eumetazoans. Furthermore, we have identified major eumetazoan cell-adhesion gene families, including those that encode cell-surface receptors, cytoplasmic linkers, and extracellular-matrix proteins. We found that approximately 50% of genes differentially expressed during postlarval development are also expressed in regeneration. The *S. ciliatum* dissociated cells thus deploy a similar set of genes as their embryonic and larval development to assist regeneration in cell culture. This has important implications for using the *S. ciliatum* regeneration system to model the evolution of multicellularity. In the current study our sampling was aimed to dissect the early cellular events in regeneration, an additional high throughput studies over multiple time points can address deeper questions such as somatic cell trajectory during regeneration.

## Methods

### Animal collection and cell culture

Adult *S. ciliatum* specimens were collected in the vicinity of Plymouth, Devon, UK. All specimens were at the stage of the post reproduction growth and did not contain any reproductive elements. Prior to experimentation, animals were maintained in the aquarium with filtered seawater (FSW) no longer than 2-4 hr. Each *S. ciliatum* was cut into small pieces by using a scalpel and micro scissors, then mechanically passed through 40 µm nylon mesh (Fisherbrand™) into a petri dish (35 mm). The dissociated cells were suspended in FSW and pipetted multiple times to allow maximum dissociation. The dissociated cells from each individual were distributed into six-well plates. The number of cells in the plates ranged from 1.2×10^6^ -1.5×10^7^. Cell cultures were maintained at 15°C in dark, every 24 hr the culture medium was exchanged with FSW. The cell culture was observed daily, and bright-field images were collected. Initially, when attempting to follow published protocols [3-5], we were unsuccessful with the *S. ciliatum* regeneration, the majority of the cell aggregates were unable to progress to functional juveniles. Hence, we modified the protocol by fractionating the dissociated cells through centrifugation. We followed similar instructions until cell dissociation, and then we centrifuged the dissociated cell at 1.2 rpm for 2 min. The supernatant was collected into a new 1.5 ml tube and re-centrifuged at 2.5 rpm for 2 min. The cell pellets from both steps were suspended in FSW and distributed individually in 6 well plates. The cells pelleted at 2.5 rpm have a high regeneration ability in comparison to the cells pelleted at 1.2 rpm. In the current study, we used the aggregates collected from 2.5 rpm wells. Apart from standardizing the protocol, the regeneration experiment was performed 7 times independently and 8 to 12 specimens were used in each experiment. For morphological analysis, the samples were collected on days post dissociation 1, 2, 3, 4, 5, 6, 8, 10, 12, 14, 18, 21 and 24. 10-20 regenerative structures were collected per replicate.

### Whole-mount immunofluorescence

The specimens were fixed for 1 hr at 4°C on a roller with 0.05% glutaraldehyde and 4% paraformaldehyde (PFA) in PBS buffer. After fixation, the samples were washed 5 times with PBST (1× PBS, 0.05% (vol/vol) Tween-20) for 5 min. For long term storage, the specimens were dehydrated in a series of ethanol dilutions and stored in 70% ethanol at 4°C. For whole-mount immunofluorescence, the samples were permeabilized in Dent’s fixative (80% methanol, 20% DMSO) for 15 minutes and followed by rehydration through series of methanol dilutions (70%, 50% and 25% methanol) for 5 minutes per dilution. The samples were washed with PBST for 3 × 10 min, then blocked in 5% BSA in PBST for 1 hr at RT. Primary antibody (1:500 dilution, mouse Anti-α-Tubulin Cat # T9026, Sigma-Aldrich) incubation was performed in a blocking solution (1% BSA in PBST) for 24-36 hr at 4°C. The samples were washed with PBST for 5 × 5 min, after which samples were incubated with secondary antibodies (1:250 dilution; Goat anti-Mouse IgG Alexa Fluor 594 Cat # A-11032, ThermoFisher) diluted in blocking solution for overnight at 4°C. Then, the samples were washed with PBST for 5 × 10 min and clarified following iDISCO protocol [45]. Imaging was performed on Leica TCS SP8 DLS and Leica DMi8 confocal microscopes.

### Fixation of regenerating specimens for Scanning Electron Microscopy (SEM)

The specimens were fixed overnight at 4°C by 2.5% glutaraldehyde in PBS buffer, then rinsed thrice with PBS buffer for 10 min. The specimens were postfixed in 1% osmium tetroxide in PBS buffer at room temperature. After post-fixation, the specimens were dehydrated in a series of ethanol dilutions until 100% ethanol and absolute ethanol was gradually exchanged with hexamethyldisilazane (HMDS). After the final 100% HMDS solution step, the specimens were left in a fume hood overnight to allow HMDS to evaporate. After overnight drying, the samples were mounted on aluminum stabs and gold coated. SEM imaging was done on JEOL 6610 LV microscope.

### RNA sequencing and differential gene expression

The regeneration is semi-synchronous among individuals, even though the variation is considerably mild (generally varies ±1 day). Keeping this in mind we collected the regenerating samples principally based on morphological signatures apart from the day 1 and 2 samples. For RNA isolation, the early regenerating samples were collected at days 1 and 2. For later stages, the samples were primarily collected based on the morphological state extending between day 3-4, day 6-8, day 10-14, day 16-18 (PJUV) and day 21-24 (JUV). Each sample was pooled from a minimum of three individuals and also included samples from different batches as detailed in table 1, 1^st^ column. The samples were carefully collected using a pipette and excess media was removed before snap freezing in liquid nitrogen and stored at -80°C until further processing. Due to the sheer size of primmorphs at early stages (day 1 and 2) samples collected from multiple biological replicates were combined to acquire an adequate amount of RNA for sequencing. For the later stages, only 3 biological replicates were combined into one sample during RNA isolation. Total RNA was isolated using the TRI Reagent® according to the manufacturer’s protocol. RNA quality was assessed using Agilent RNA 6000 Nano Kit on Agilent 2100 Bioanalyzer (Agilent, USA), and only samples with RNA integrity number (RIN) ≥ 8.0 were considered for sequencing. Sample library preparation for RNA sequencing was accomplished using the SENSE mRNA-Seq Library Prep Kit (Lexogen GmbH). Before sequencing, the libraries were pre-assessed by Agilent High Sensitivity DNA Kit (Agilent, USA) and quantified using Qubit™ 1X dsDNA HS Assay Kit (Invitrogen™). The sequencing was outsourced (GENEWIZ Illumina NovaSeq/HiSeq 2×150 bp sequencing), generating a 20 million paired-end reads per replicate. Raw data was deposited at NCBI GEO submission GSE149471. After de-multiplexing and filtering high-quality sequencing reads, the adapter contamination was removed by using Trimmomatic v0.36 [46]. The quality of the reads was verified using FastQC [47]. Processed reads from each sample were mapped to the *S. ciliatum* genome (indexed bowtie2 [48]) by using HISAT2 [49]. Next, the mapped reads were passed to StringTie [50] for transcript assembly. After initial assembly, all assembled transcripts from regeneration and development were merged by using StringTie’s merge module, which merges all the gene structures found in any of the samples. The BAM file and GFT file generated by StringTie’s merge were fed into featureCounts [51] to extract the reads counts per transcript. Differential expression analyses were performed using DESeq2 (Galaxy Version 2.11.40.6) [52]. Gene models that did not have >10 counts in at least 25% of the samples were excluded. The principal component analysis (PCA), hierarchical clustering (HC), and heat maps were generated using the R package in R-studio (Version 1.2.5019). For functional annotation, the transcripts were extracted from the GTF annotation file by gffread utility and the LongOrfs were acquired using TransDecoder. We used blastp [53] with the default curated gathering threshold to predict the protein orthologues against the Uniprot database. The Gene Ontology (GO) term enrichment was performed using gene annotation tools including PANTHER™ Classification System [54] and DAVID [55].

## Acknowledgments

The authors would like to thank John Bishop and Christine Wood for their support in acquiring *S. ciliatum* specimens. We thank Glenn Harper and the team at the Plymouth Electron Microscopy Lab for their assistance during the imaging. We thank Ro Allen for his help with R statistics. This work was supported by the Anne Warner endowed Fellowship through the Marine Biological Association of the UK.

---

**Supplementary File 1: 4D and Z-stack of imaging of primmorphs after 24 hrs post dissociation**.

**Supplementary File 2: 4D and Z-stack of imaging of primmorphs 2 days post dissociation**.

**Supplementary File 3: 4D and Z-stack of imaging of primmorphs 3 days post dissociation**.

**Supplementary File 4: 4D and Z-stack of imaging of structures with ciliated chambers 8 days post dissociation**.

**Supplementary File 5: 4D and Z-stack of imaging of regenerating structure at 12 days post dissociation**. Defined choanoderm with a single layer of choanocytes surrounding the atrium. Note that ostium and osculum are not formed yet.

**Supplementary File 6: 4D and Z-stack of imaging of pre-juvenile 18 days post dissociation**. The ostium and osculum are formed.

**Supplementary File 7: 4D and Z-stack of imaging of juvenile 24 days post dissociation**.

**Supplementary File 8:**
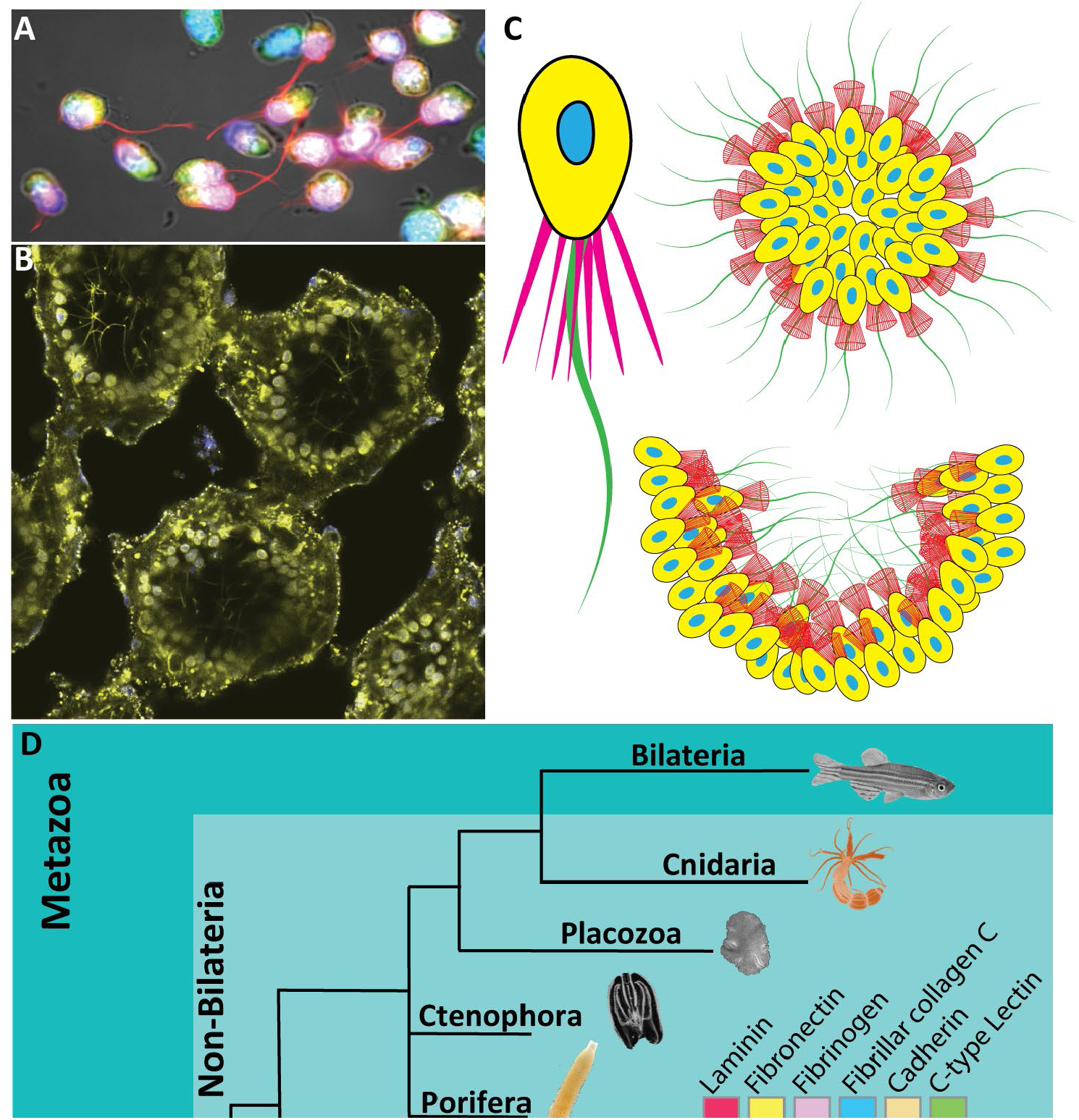
Differentially expressed genes associated with cell death. **(A)** Heatmap shows the relative expression of genes associated with cell death (apoptosis signaling pathway, P38 MAPK, and response to oxidative stress). The selected list of genes is significantly DE across the course of *S. ciliatum* regeneration. Note the large changes in gene expression at Day 1 and the gradual decrease in the expression later on.

